## Supplementary Figures for "A coordinated transcriptional switching network mediates antigenic variation of human malaria parasites"

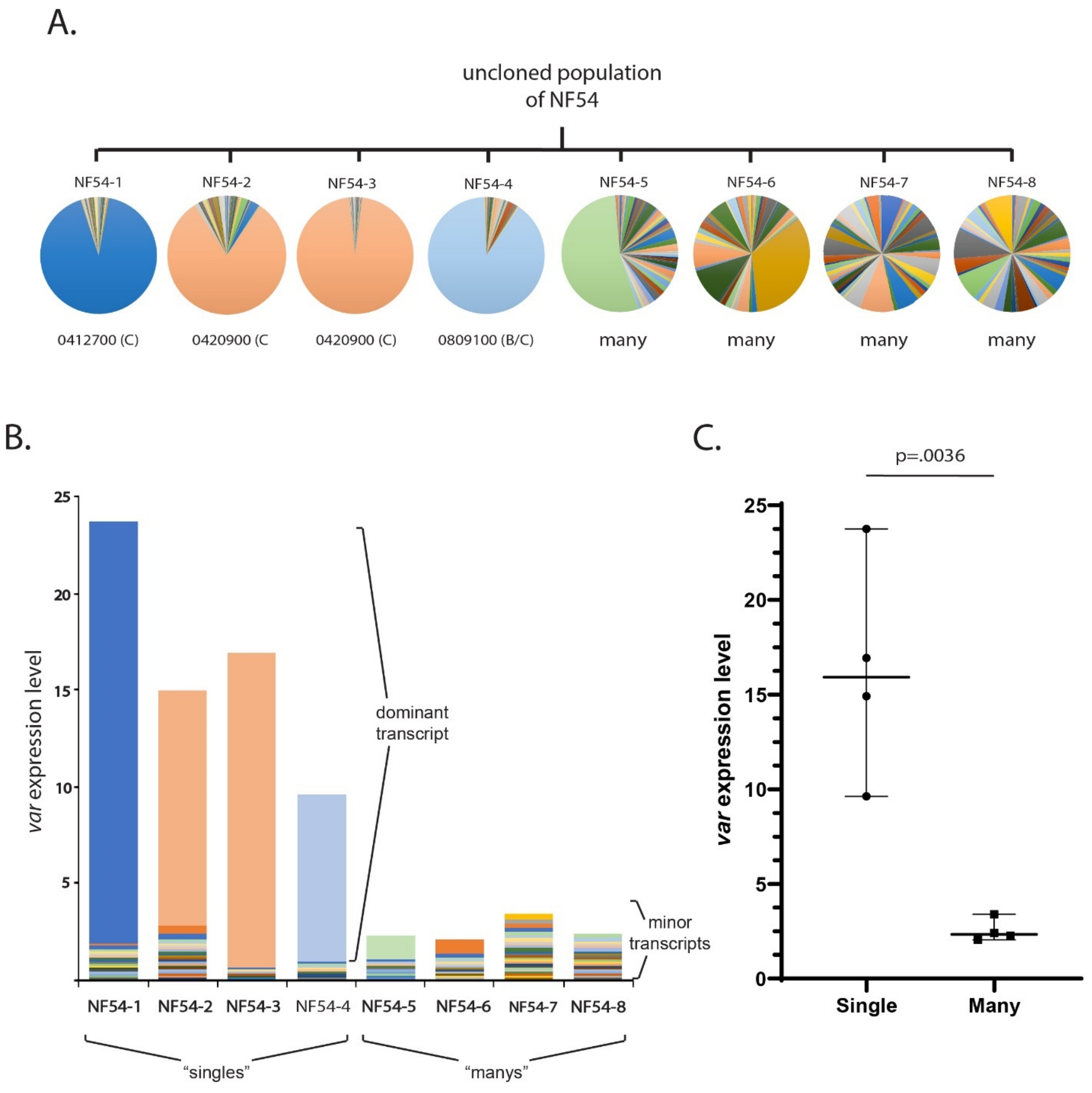


Supplementary Figure 1: Examination of *var* gene expression profiles of recent clones of NF54. A) Cloning by limiting dilution was performed using an uncloned, heterogenous population of the NF54 isolate. 8 individual clonal populations were generated and allowed to expand for 5 weeks, at which point they were synchronized and RNA isolated from ring-stages. *var* gene expression profiles were generated and displayed as pie charts with the dominantly expressed *var* gene indicated below the chart for clones 1-4. B) Total *var* expression levels for all eight clones shown in A, with transcripts for each *var* gene shown in a different color. Note that for clones NF54-5 and NF54-6, one *var* gene is more highly expressed than the rest of the family (as can be observed in the pie charts in A, however the overall *var* expression level was very low, and therefore these clones were classified as “manys”. C) Total *var* expression levels as determined by qRT-PCR for all eight NF54 subclones. The median +/- 95% confidence interval is shown, and an unpaired t-test indicates a p value of .0036.

A.

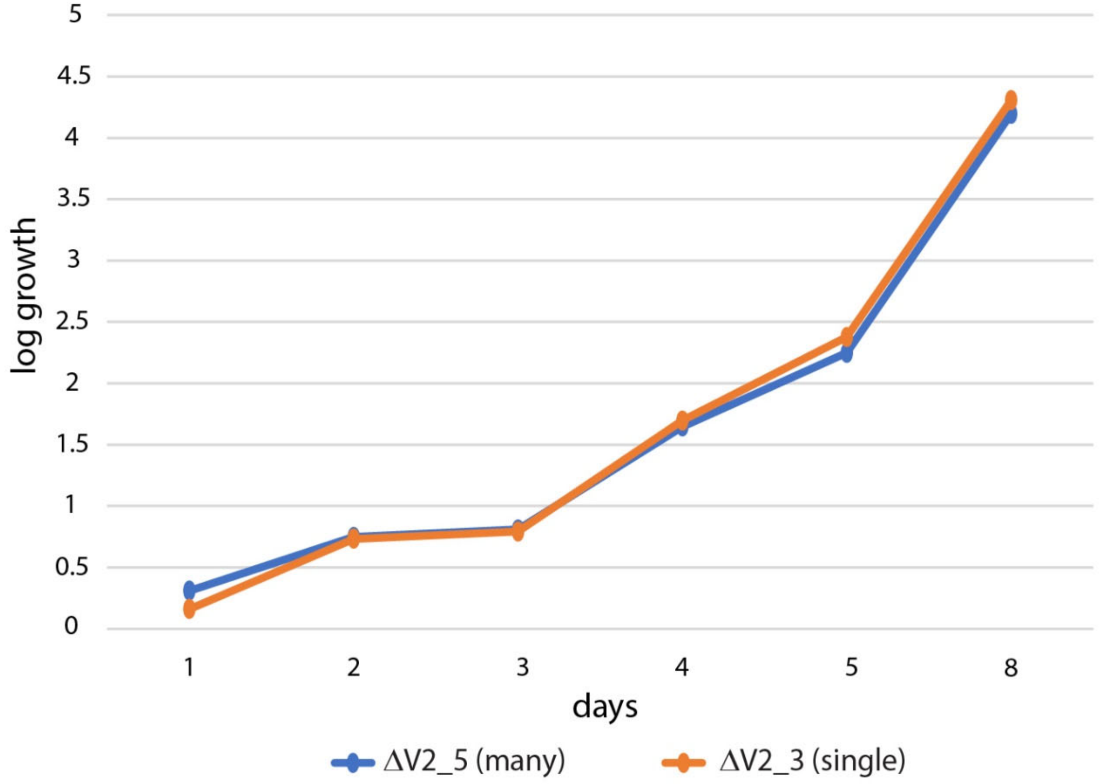

5
4.5
4
3.5
3
2.5
2
1.5
1
0.5
0
log growth
1
2
3
4
5
8
days
ΔV2_5 (many)
ΔV2_3 (single)


B.

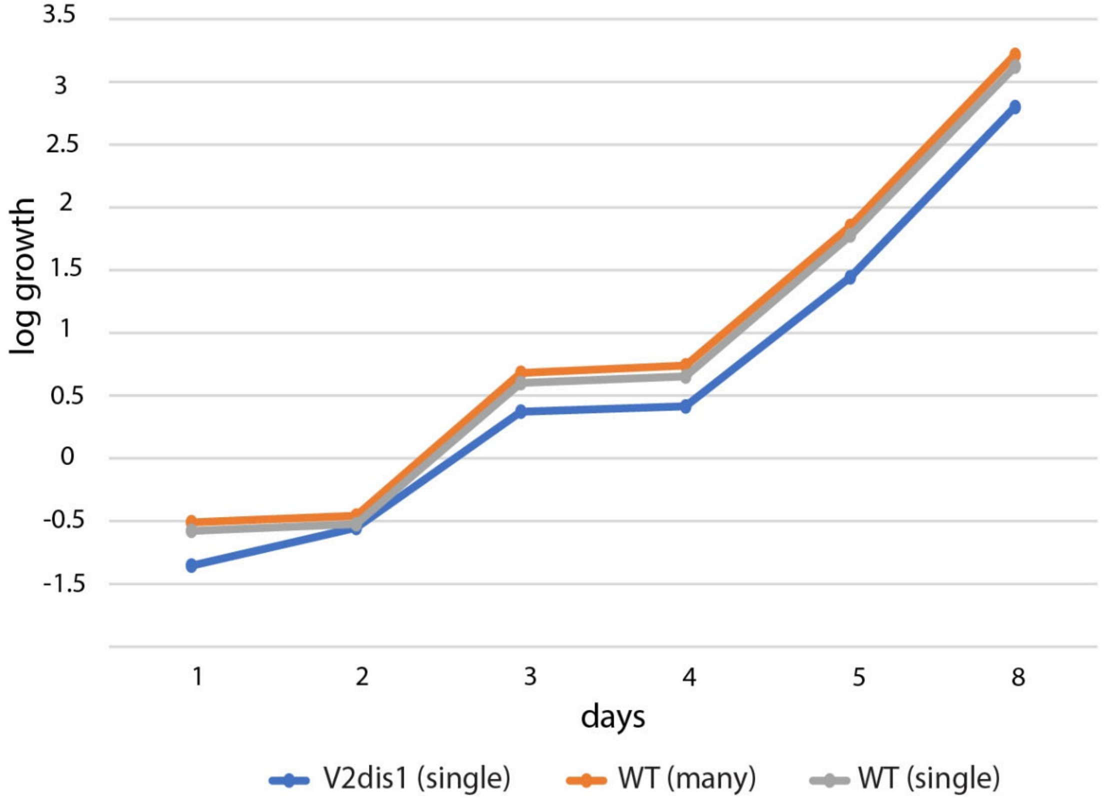

3.5
3
2.5
2
1.5
1
0.5
0
-0.5
-1.5
log growth
1
2
3
4
5
8
days
V2dis1 (single)
WT (many)
WT (single)

Supplementary Figure 2: Growth assays of parasites displaying the “single” and “many” *var* expression phenotypes. Parasitemias were determined daily by flow cytometry and cultures were diluted by 10 fold whenever the parasitemia reached 0.5%. The daily parasitemia was multiplied by the exponential dilution factor and plotted as log growth over time. A) Comparison of two clonal lines in which *var2csa* has been deleted. ΔV2_5 (blue) displays the “many” phenotype while ΔV2_3 (orange) displays the “single” expression profile. B) Comparison of a *var2csa*-mutant line (V2dis1) displaying the “single” phenotype with 2 wildtype lines, one displaying the many phenotype (orange) and one expressing a single dominant *var* gene (grey).
